# The metabolic benefits of thermogenic stimulation are preserved in aging

**DOI:** 10.1101/2024.07.01.601572

**Authors:** Duraipandy Natarajan, Bhuvana Plakkot, Kritika Tiwari, Shoba Ekambaram, Weidong Wang, Michael Rudolph, Mahmoud A. Mohammad, Shaji K. Chacko, Madhan Subramanian, Stefano Tarantini, Andriy Yabluchanskiy, Zoltan Ungvari, Anna Csiszar, Priya Balasubramanian

## Abstract

Adipose thermogenesis has been actively investigated as a therapeutic target for improving metabolic dysfunction in obesity. However, its applicability to middle-aged and older populations, which bear the highest obesity prevalence in the US (approximately 40%), remains uncertain due to age-related decline in thermogenic responses. In this study, we investigated the effects of chronic thermogenic stimulation using the β3-adrenergic (AR) agonist CL316,243 (CL) on systemic metabolism and adipose function in aged (18-month-old) C57BL/6JN mice. Sustained β3-AR treatment resulted in reduced fat mass, increased energy expenditure, increased fatty acid oxidation and mitochondrial activity in adipose depots, improved glucose homeostasis, and a favorable adipokine profile. At the cellular level, CL treatment increased uncoupling protein 1 (UCP1)-dependent thermogenesis in brown adipose tissue (BAT). However, in white adipose tissue (WAT) depots, CL treatment increased glycerol and lipid de novo lipogenesis (DNL) and turnover suggesting the activation of the futile substrate cycle of lipolysis and reesterification in a UCP1-independent manner. Increased lipid turnover was also associated with the simultaneous upregulation of proteins involved in glycerol metabolism, fatty acid oxidation, and reesterification in WAT. Further, a dose-dependent impact of CL treatment on inflammation was observed, particularly in subcutaneous WAT, suggesting a potential mismatch between fatty acid supply and oxidation. These findings indicate that chronic β3-AR stimulation activates distinct cellular mechanisms that increase energy expenditure in BAT and WAT to improve systemic metabolism in aged mice. Our study provides foundational evidence for targeting adipose thermogenesis to improve age-related metabolic dysfunction.

## Introduction

Aging is accompanied by detrimental changes in adipose tissue, characterized by immune cell infiltration, preadipocyte senescence, reduced sympathetic input, increased fibrosis, and altered adipokine secretion^1,2^. These pathological alterations in the adipose tissue contribute to whole-body metabolic dysfunction through ectopic lipid deposition, insulin resistance, and low-grade chronic inflammation, all of which are implicated in accelerating systemic aging^3^. Notably, metabolic dysfunction in middle age precedes the onset of several age-related diseases^4–8^, underscoring the potential of interventions targeting adipose tissue metabolism to extend the healthspan of aging individuals.

The discovery of metabolically active brown adipose tissue (BAT) in adult humans over 15 years ago^9–11^ sparked renewed interest in adipose thermogenesis as a means to combat metabolic dysfunction in obesity. Evidence from preclinical and clinical studies suggests that thermogenic stimulation through cold exposure or β3-adrenergic (AR) agonists enhances cellular energy expenditure by catabolizing glucose, free fatty acids (FA), and branched-chain amino acids in activated brown and beige adipocytes (brown-like adipocytes arising within white adipose tissue (WAT) through a process called browning or beiging)^12–14^. At the molecular level, uncoupling protein 1 (UCP1), expressed in the inner mitochondrial membranes of brown and beige adipocytes, mediates the thermogenic response by promoting heat generation through uncoupling the electron transport chain (ETC) activity from ATP synthesis^15^. Importantly, increased fuel mobilization and utilization in activated brown and beige adipocytes containing elevated UCP1 led to improvements in glucose tolerance and insulin sensitivity^16,17^. Adrenergic stimulation increases energy expenditure in both BAT and WAT; however, UCP1 expression varies greatly among adipose depots (with BAT showing the highest expression, followed by inguinal or subcutaneous WAT (iWAT), and then epididymal or visceral WAT (eWAT)). Similar increases in oxidative metabolism despite varying levels of UCP1 between adipose depots suggest that cellular mechanisms independent of UCP1 might also contribute to energy expenditure. In concordance with this idea, recent studies have highlighted the existence of noncanonical thermogenic mechanisms that occur in beige adipocytes independent of UCP1 wherein energy expenditure is primarily mediated through the ATP-consuming futile cycling of metabolic substrates, including triglyceride (TG)-FA, calcium, and creatine^18–22^.

Although obesity and aging share several metabolic phenotypes associated with dysfunctional adipose tissue^23^, the therapeutic potential of thermogenesis in aged individuals remains largely unexplored^24^. This knowledge gap can be partly attributed to reported losses in classical BAT tissue and impaired beige adipocyte formation in WAT in response to cold or β3-AR stimulation during aging^25,26^. Nevertheless, cold stimulation has been used in a majority of studies to investigate the thermogenic response in aged animals^27, 28,28^. Given that the sympathetic output to the adipose tissue declines with age^29^, the maximum potential of thermogenesis may not be attained with cold stimulation. This limitation can be avoided by direct activation of β3-AR receptors via pharmacological agents and might be more appropriate for assessing thermogenesis in aged animals because β3-AR stimulation bypasses the sympathetic nervous system. A major question in the aging field is whether thermogenic responses elicited in aged animals are sufficient to improve age-related metabolic outcomes, especially given the high incidence of obesity and metabolic disease among the aging population. Furthermore, whether UCP1-independent thermogenic mechanisms are preserved during aging has not been established in mouse models. Accordingly, we tested the hypothesis that UCP1-independent thermogenic mechanisms following chronic β3-AR treatment are preserved during aging and lead to improvements in systemic metabolic outcomes, adipose function, and inflammation of late middle-aged mice. A better understanding of UCP1-independent thermogenic mechanisms during aging could provide new therapeutic options for the treatment of metabolic dysfunction, particularly in elderly populations with compromised UCP1-dependent thermogenesis.

## Materials and Methods

### Animals and treatment

Aged C57BL/6JN male and female mice (18 months old) were obtained from an aging colony maintained by the National Institute on Aging, Charles River Laboratories (Wilmington, MA, USA). The animals were housed in a conventional animal housing facility with a 12:12-hour light-dark cycle at the University of Oklahoma Health Sciences Center. They were fed a standard chow diet (PicoLab Rodent Diet 5053) ad libitum with continuous access to water and enrichment. The animals underwent sham surgery (controls) or were implanted with osmotic minipumps filled with β3-AR agonist (CL 316,243 [CL]); R&D Systems-Cat. No. 1499/50; 0.75 nmol/h)^20^ to enable continuous infusion for 4 weeks. Body composition was measured before and after treatment using an EchoMRI Body Composition Analyzer (Houston, TX, USA). Mice were euthanized in a fed state at the end of 6 weeks of CL treatment, and serum, eWAT, iWAT, and BAT tissues were collected, followed by storing at −80 °C for RNA and protein analysis or fixing in 10% formalin for paraffin embedding. Fresh tissues were collected for 2,3,4-Triphenyltetrazolium (TTC) staining, as described below. All animal experiments were approved by the Institutional Animal Care and Use Committee of the University of Oklahoma Health Sciences Center.

### Indirect calorimetric assay

Oxygen consumption (VO_2_), carbon dioxide production (VCO_2_), energy expenditure (EE), food intake, and activity were assessed using a Promethion Core Monitoring System (Sable Systems, Las Vegas, NV, USA). Briefly, the animals were individually housed in chambers and allowed to acclimate for 24 hours. After acclimatization, metabolic parameters were recorded every 5 min for 3 d. VO_2_ and EE values were normalized to the body weight of the animals before analysis. The respiratory exchange ratio (RER) was calculated by dividing CO_2_ produced by O_2_ consumed.

### *In situ* mitochondrial ETC activity analysis

We evaluated mitochondrial ETC activity in freshly isolated adipose tissues by staining with the redox dye TTC as described previously^30^. Briefly, adipose tissue (50–75 mg) was minced into small pieces and incubated at 37 °C in 1% TTC in Phosphate-buffered saline for 30 min. Insoluble red formazan, which is the reduced form of TTC, was extracted in 100% isopropanol and incubated overnight at room temperature (RT). The absorbance of isopropanol was measured at 485 nm, and the OD was normalized to the tissue weight.

### Glucose tolerance test

After overnight fasting, blood was collected to measure fasting glucose and insulin levels. Mice were then given a bolus of glucose (2 g/kg body weight) in saline delivered via intraperitoneal injections of 250–500 μl of liquid, depending on the size of the animal. Blood glucose levels were monitored with a handheld glucometer (OneTouch) using fresh blood collected from the tail nick at the end of the tail at 0 (before glucose injection), 15, 30, 60, and 120 min after injection. HOMA-IR (homeostasis model assessment of insulin resistance) index was calculated as [fasting serum glucose×fasting serum insulin/22.5].

### Insulin sensitivity assay

Mice fasted for 4-6 hrs were given insulin (2mU/g) intraperitoneally. After 15 minutes of insulin injection, the mice were sacrificed by cervical dislocation, and the liver and gastrocnemius muscle were collected to assess protein levels of pAKT/AKT by western blotting.

### Western blotting

Tissue samples were homogenized in ice-cold radioimmunoprecipitation assay lysis buffer (Millipore Sigma, #R0278) containing a Halt protease and phosphatase inhibitor cocktail (Thermo Scientific, #PI78440). The lysates were placed in a rotating chamber in a cold room for 1 h and centrifuged at 16,000 g for 10 min. A clear layer was collected at the bottom, leaving a lipid layer at the top. Protein concentrations were assessed using the Pierce BCA Protein Assay Kit (Thermo Scientific, #23227). Equal amounts of protein (30 µg) were separated on a 4–20% SDS gel under reduced and denatured conditions, followed by wet transfer to a PVDF membrane (Bio-Rad). The transfer was confirmed using Ponceau staining and imaging, followed by washing twice with Tris-buffered saline with 1% tween 20 (TBST) for 5 min each. After blocking for 1 h with 5% non-fat milk at RT, the membranes were incubated overnight in a cold room with the following primary antibodies: anti-AKT (1:1000, Cell Signaling #9272S), anti-pAKT (1:1000, Cell signaling #9271S), anti-UCP1 (1:1000, R&D Systems #MAB6158), anti-glycerol kinase (GYK, 1:1000, Abcam #ab126599), anti-phosphoenolpyruvate carboxykinase (PEPCK, 1:1000, Abcam #ab70358), anti-acyl-coA synthetase 1 (ACSL1, 1:1000, Cell signaling #4047S), anti-diacylglycerol O-acyltransferase 1 (DGAT1, 1:1000, Novus Biologicals #NBP1-71701) and anti-medium-chain acyl-coenzyme A dehydrogenase (MCAD, 1:1000, Santa Cruz #sc365108). Following washing with TBST thrice, the membranes were incubated with the respective horseradish peroxidase-conjugated secondary antibodies (Abcam #ab205719 and ab6721) for 1 h at RT. Following washing, the bands were developed using Pierce SuperSignal West Pico Plus chemiluminescent substrate, and the signals were detected using the ChemDoc imaging system. Densitometry was performed using Fiji ImageJ software, and the values were normalized to Ponceau staining.

### In vivo evaluation of glycerol and lipid synthesis using stable isotopes

During the last 10 days of CL treatment, mice were enriched with deuterium-labeled water (D_2_O) for isotope tracer experiments to measure newly synthesized triglycerides and fatty acids. Aged mice treated with saline or CL were administered a bolus of labeled water by intraperitoneal injections (isotonic saline containing >99% ^2^H_2_O @ 25µl/g of lean body mass). After the injections, the mice were maintained on 5% ^2^H_2_O in drinking water to maintain a steady state labeling of body water for the next 10 days. Urine was collected 24 hrs after the ip injections and the serum and fat tissues were collected at the end of the 10 days to confirm label incorporation. Newly synthesized TG was calculated from the incorporation of ^2^H into TG-derived glycerol, and newly synthesized palmitate was determined from the incorporation of ^2^H into TG-derived palmitate. All measurements were made in the Stable Isotope Core Laboratory of the USDA/Children’s Nutrition Research Center, Baylor College of Medicine, Houston, TX. Briefly, eWAT& iWAT TGs were hydrolyzed with 2.0 N KOH containing 50% ethanol at 110°C for 2 hours, and the incorporation of ^2^H into glycerol and palmitate was determined by gas chromatography-mass spectrometry (GC-MS) following derivatization to glyceryl triacetate ^31^ and palmitate pentaflurobenzylbromide derivative^32^. 2H2O enrichment was measured using Isotopic Ratio Mass Spectrometry as described previously (IRMS)^32^.

### Fatty acid oxidation assay

Fatty acid β-oxidation activity was assessed in lysates prepared from frozen adipose tissue using a commercial kit (Assay Genie). This assay is based on the oxidation of octanoyl-CoA, which is coupled with NADH-dependent reduction, and the formation of a formazan product that can be measured at an absorption wavelength of 492 nm. Values were normalized to the total protein content of the samples and expressed as units/µg of protein.

### Milliplex assays for cytokine and adipokine analysis

Protein lysates from the adipose tissue and serum samples were analyzed for inflammatory markers (Millipore Sigma #MCYTOMAG-70K-PMX) and adipokines (Millipore Sigma #MADMAG-71K-07) using Milliplex kits. The values from protein lysates were normalized to the total protein content in each sample and expressed as pg/µg of protein.

### Enzyme-linked immunosorbent assay (ELISA) and biochemical assays

Serum adiponectin levels were measured using Quantikine ELISA kits (#MRP300; R&D Systems). Triglyceride and free fatty acid levels were assessed using colorimetric kits (Cayman Chemicals #10010303 and Sigma-Aldrich #MAK044), respectively.

### Histological and immunohistochemical assays

The fat tissues were fixed in 4% PFA, paraffin-embedded, sectioned at 5 µm thickness, and mounted on positively charged slides. Slides were transferred to Leica Bond RX for dewaxing and then treated for target retrieval at 100 °C for 20 min in a retrieval solution at pH 6.0. The sections were incubated with 5% goat serum (ThermoFisher Scientific #01-6201) for 30 min. Endogenous peroxidase was blocked using a peroxidase-blocking reagent, followed by incubation with UCP1 (Abcam #ab10983) for 60 min and then with a Poly-HRP IgG secondary antibody. Detection was done using 3,3′-diaminobenzidine tetrahydrochloride (DAB) as a chromogen and counterstaining with hematoxylin. Slides were dehydrated (Leica ST5020) and mounted (Leica MM24). Digital images were acquired using an AxioScan microscope slide scanner (Zeiss, Oberkochen, Germany). Adipocyte area measurements were performed on hematoxylin and eosin (H&E)-stained slides using the AdipoSoft plug-in in the ImageJ software.

### Statistical analysis

Statistical analyses were performed using GraphPad Prism 9.3.1 (GraphPad Software, San Diego, CA, USA), and the data are expressed as mean ± standard error of the mean (SEM). The data were analyzed using a two-tailed unpaired Student’s t-test. The level of statistical significance was set at p < 0.05.

## Results

### Chronic β3-AR stimulation promotes fat loss and improves insulin sensitivity in aged mice

To investigate the metabolic response to prolonged β3-AR stimulation in aged mice, 18-month-old C57BL/6J mice were treated with CL for 4 weeks via osmotic minipump infusion at 0.75 nmol/hr. CL treatment significantly decreased whole animal fat mass and increased the lean mass as a function of total body weight, which remained unchanged, relative to controls (Fig. 1A-C). Fat loss was also reflected in the tissue weights of individual fat pads, where there was a significant reduction in eWAT and iWAT mass in CL-treated mice. However, we did not observe any changes in the BAT or liver mass following CL treatment in aged mice (Supp. Fig. 1A-D). Fasting glucose, fasting insulin, and the insulin resistance measure homeostatic model assessment for insulin resistance (HOMA-IR) were significantly decreased following CL treatment in aged mice (Fig. 1D-F). Consistent with these findings, CL-treated animals also showed improved glucose tolerance, as indicated by a reduced area under the curve (AUC) (Fig. 1G-H). Regarding adipokines, CL treatment increased circulating adiponectin levels; however, no differences were observed in leptin levels (Fig. 1I). Serum FA and triglyceride levels were also not different between the control and CL groups (Suppl. Fig. 1E-F). We also evaluated peripheral insulin sensitivity by measuring the phosphorylation status of AKT in the liver and gastrocnemius muscle in response to a bolus of insulin. We found significantly increased pAKT levels in both the liver and muscle of the CL-treated animals, indicating improved peripheral insulin sensitivity (Fig. 1J). These findings suggest that chronic stimulation of β3-AR improves age-related metabolic outcomes, including reductions in fat mass, improvements in glucose clearance and insulin sensitivity, and increased levels of the principle adipose-secreted adipokine, adiponectin.

**Figure 1:**
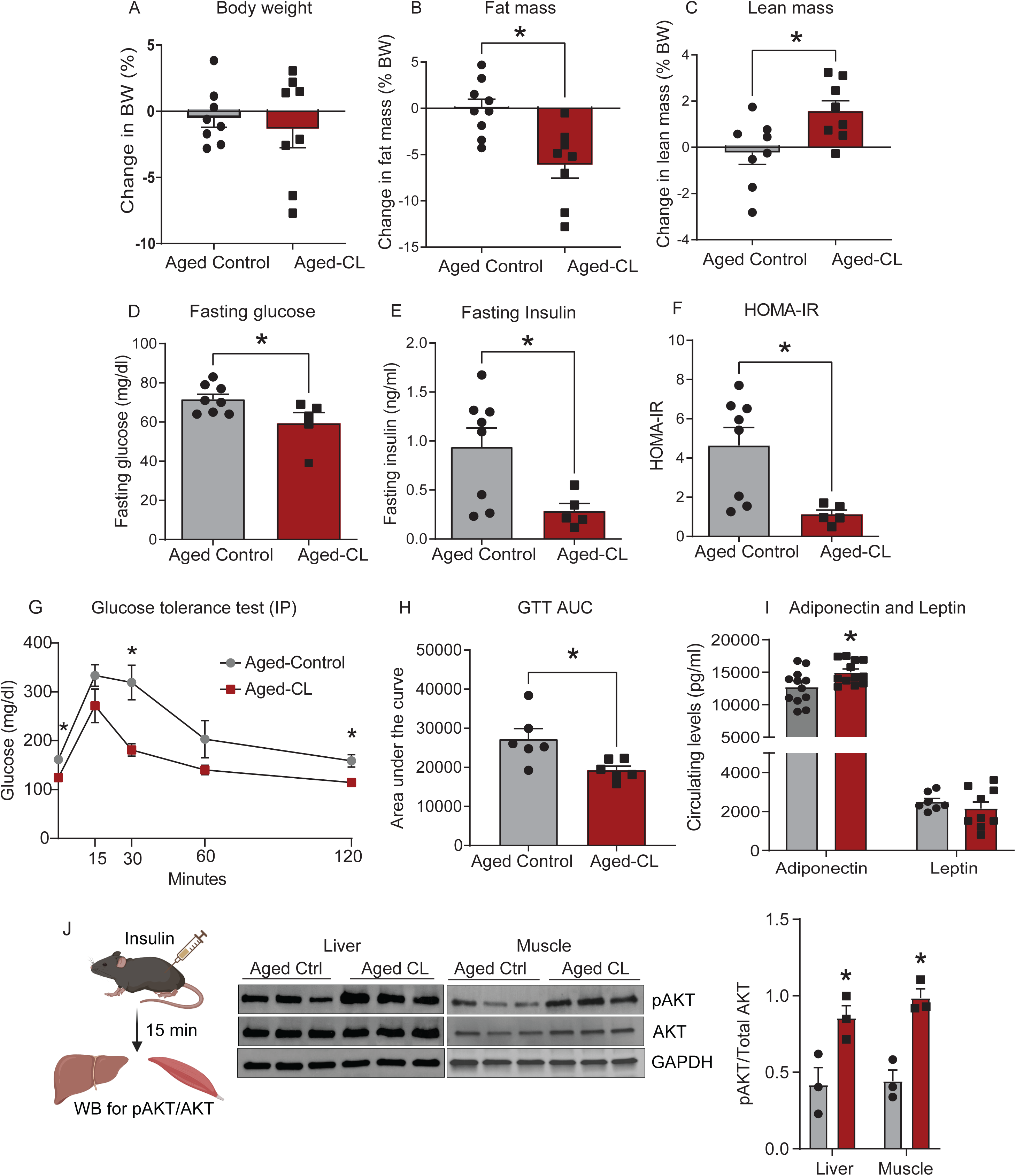
Metabolic effects of chronic β3-AR treatment in aged mice. Aged C57BL/6J mice (18 months old, both sexes) were treated with CL 316,243 (1 mg/kg BW; osmotic pump infusion for 4 weeks). Relative changes in (A-C) body weight (BW), and fat and lean masses (n = 8/group); (D-F) fasting glucose, insulin, and HOMA-IR (n = 5–8/group); (G-H) glucose values during intraperitoneal glucose tolerance test and AUC analysis (n = 6/group); I) serum FFA and TG (n = 5–12/group); and (J) adiponectin and leptin levels (aged control, n = 7–12 and aged CL, n = 9– 12). Data are shown as average ± SEM, with significance determined using Student’s t-test (*p < 0.05 between the groups).

### Chronic β3-AR stimulation increases whole-body energy expenditure and adipose fatty acid oxidation in aged mice

Sustained β3-AR activation treatment has been demonstrated to increase whole-body energy expenditure in young mice^33,34^. In the present study, we investigated whether this response is preserved in aged mice. CL treatment increased energy expenditure and oxygen consumption in aged mice in both the dark and light phases; however, the effect was more pronounced in the dark phase (active period in mice) (Fig. 2A-D). Core body temperature was also significantly increased with CL treatment in aged mice (Fig. 2E). There were no differences in ambulatory activity levels between the groups (Fig. 2F), indicating that overall metabolic rate, rather than physical activity, was the likely cause of the observed increased energy expenditure in CL-treated aged mice. Contrary to expectations, CL-treated mice did not display a preference for fat utilization, as there were no changes in RER between the groups (Fig. 2G). Additionally, we investigated whether increased metabolic activity was also observed at the level of the adipose tissue. We determined the *in situ* mitochondrial respiration of freshly isolated adipose tissue by assessing the reduction in the electron acceptor dye TTC. Chronic CL treatment significantly increased the mitochondrial ETC activity in both white and brown adipose tissue depots (Fig. 2H).

**Figure 2:**
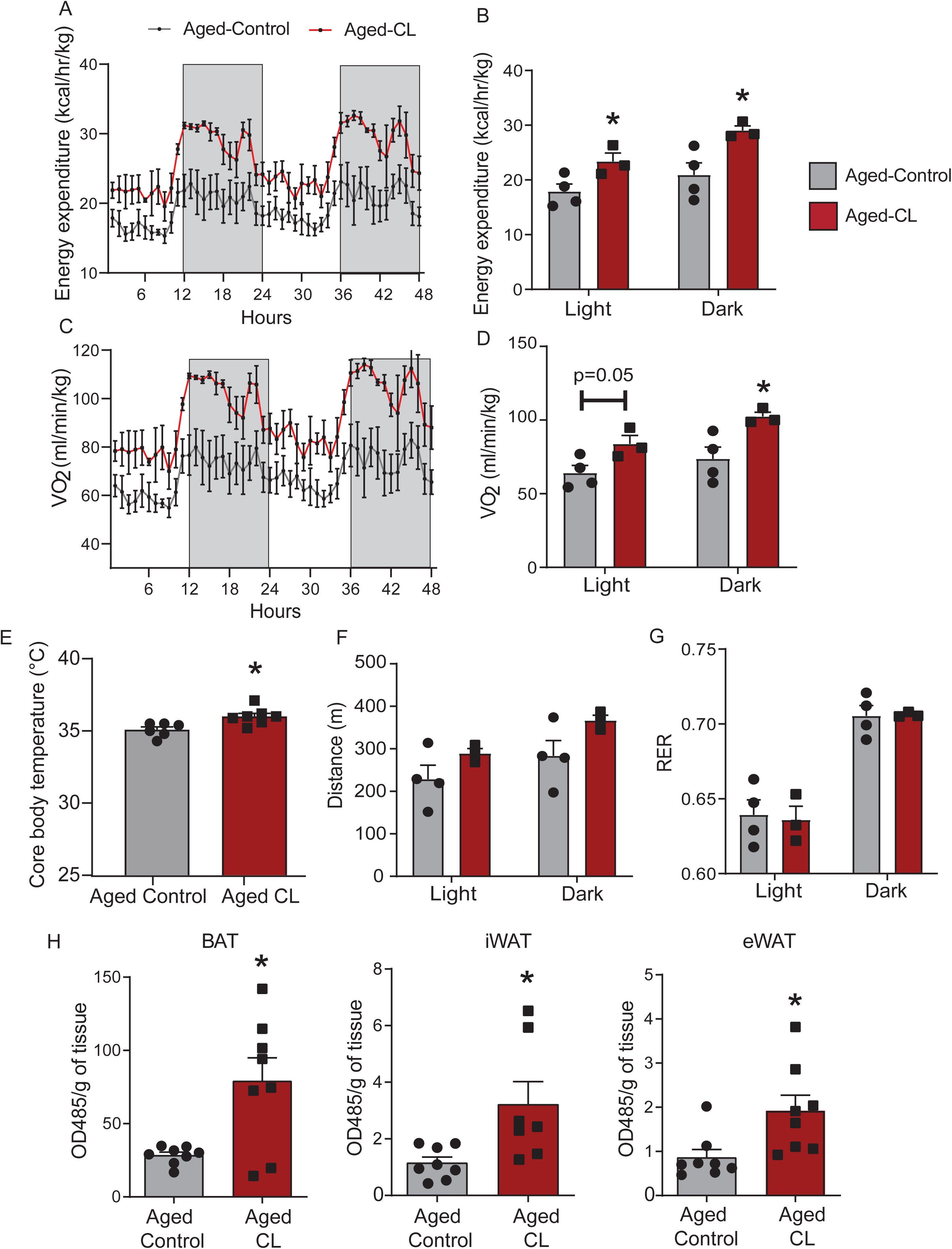
Effects of chronic β3-AR treatment on energy expenditure and mitochondrial activity in the adipose tissue depots of aged mice. Chronic CL 316,243 treatment-induced changes in (A-B) energy expenditure; (C-D) oxygen consumption; (E) Core body temperature (**֯**C); F) activity levels assessed by distance (m/mouse); and (G) RER in light and dark cycles (n = 3– 4/group; females). (H) Mitochondrial respiration assessed by measuring the reduction of the electron acceptor dye TTC in BAT, iWAT, and eWAT depots (n = 7–8/group; both sexes). Data are shown as average ± SEM, with significance determined using Student’s t-test (*p < 0.05 between the groups).

Free fatty acids released during CL-mediated lipolysis activate PPARα-mediated transcription of genes involved in fatty acid oxidation and mitochondrial electron chain activity to expand the overall oxidative capacity of adipose tissue^20,35^. Hence, we investigated whether CL treatment increases fatty acid oxidation in WAT and BAT depots. Consistent with the increased mitochondrial ETC activity shown in Fig. 2G-I, we observed increased FAO in iWAT and surprisingly not in BAT (Fig. 3A). However, FAO activity in eWAT was the lowest of all fat pads, and the values fell below the detection limit of the assay. Alternatively, we performed western blotting to assess the protein levels of Medium-chain acyl-coenzyme A dehydrogenase (MCAD), a key enzyme involved in mitochondrial fatty acid β-oxidation. MCAD protein levels in eWAT tended to increase with CL treatment (Fig. 3B). Histological analysis of H&E-stained adipose sections supported CL-induced lipid mobilization and utilization in the adipose tissue of aged mice. Both iWAT and eWAT displayed a higher ratio of smaller/larger adipocytes in CL-treated mice than in aged controls (Fig. 3C-D). These data indicate that β3-AR activation-mediated increase in energy expenditure and fatty acid oxidation are preserved in aged mice.

**Figure 3:**
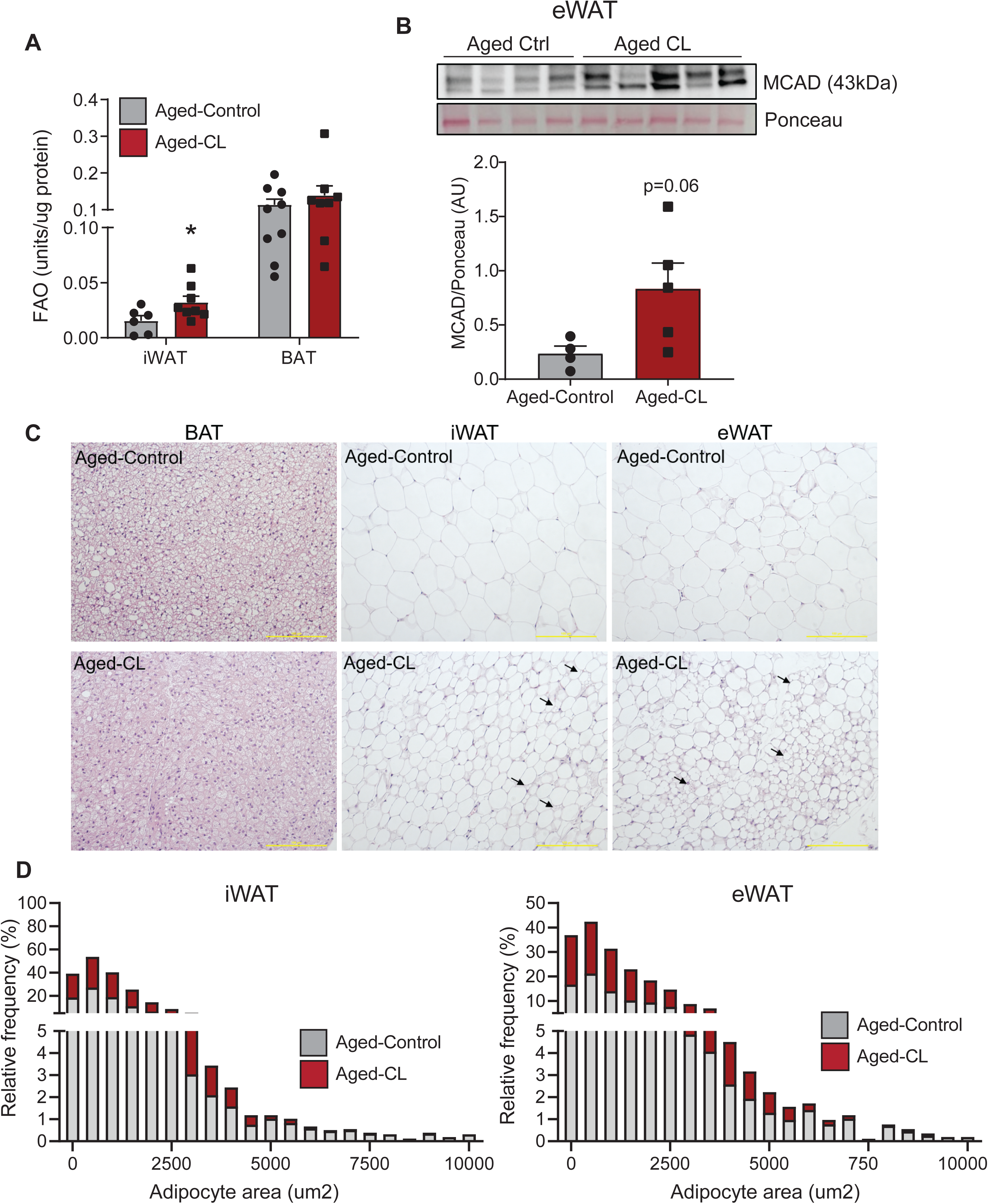
Effects of chronic β3-AR treatment on adipose fatty acid oxidation and adipocyte size distribution in aged mice. Chronic CL 316,243 treatment-induced changes in (A) fatty acid β-oxidation in iWAT and BAT depots (n = 6–9/group; both sexes) and (B) MCAD protein levels assessed using western blotting (n = 4–5/group; both sexes). (C) Representative images of H&E stained adipose sections. Black arrows indicate multilocular adipocytes. (D) Adipocyte size distribution in iWAT and eWAT. Data are shown as average ± SEM; *p < 0.05 between the groups based on the Student’s-t-test analysis.

### CL-mediated energy expenditure is mediated by UCP1 in BAT and TG-FA cycling in the WAT of aged mice

To investigate whether thermogenic mechanisms contribute to the increased mitochondrial activity in the adipose tissue of CL-treated aged mice, we first assessed UCP1 expression in various adipose depots. CL treatment significantly upregulated UCP1 protein expression in the BAT of aged mice (Fig. 4A). However, in line with prior reports^25,26^, UCP1 expression was not affected by adrenergic stimulation in the iWAT, suggesting impaired beiging with aging (Fig. 4A). We also confirmed these findings by IHC where a significant increase in multilocular UCP1+ adipocytes was observed only in the BAT but not in the WAT depots of CL -treated mice when compared to controls (Fig. 4B).

**Figure 4:**
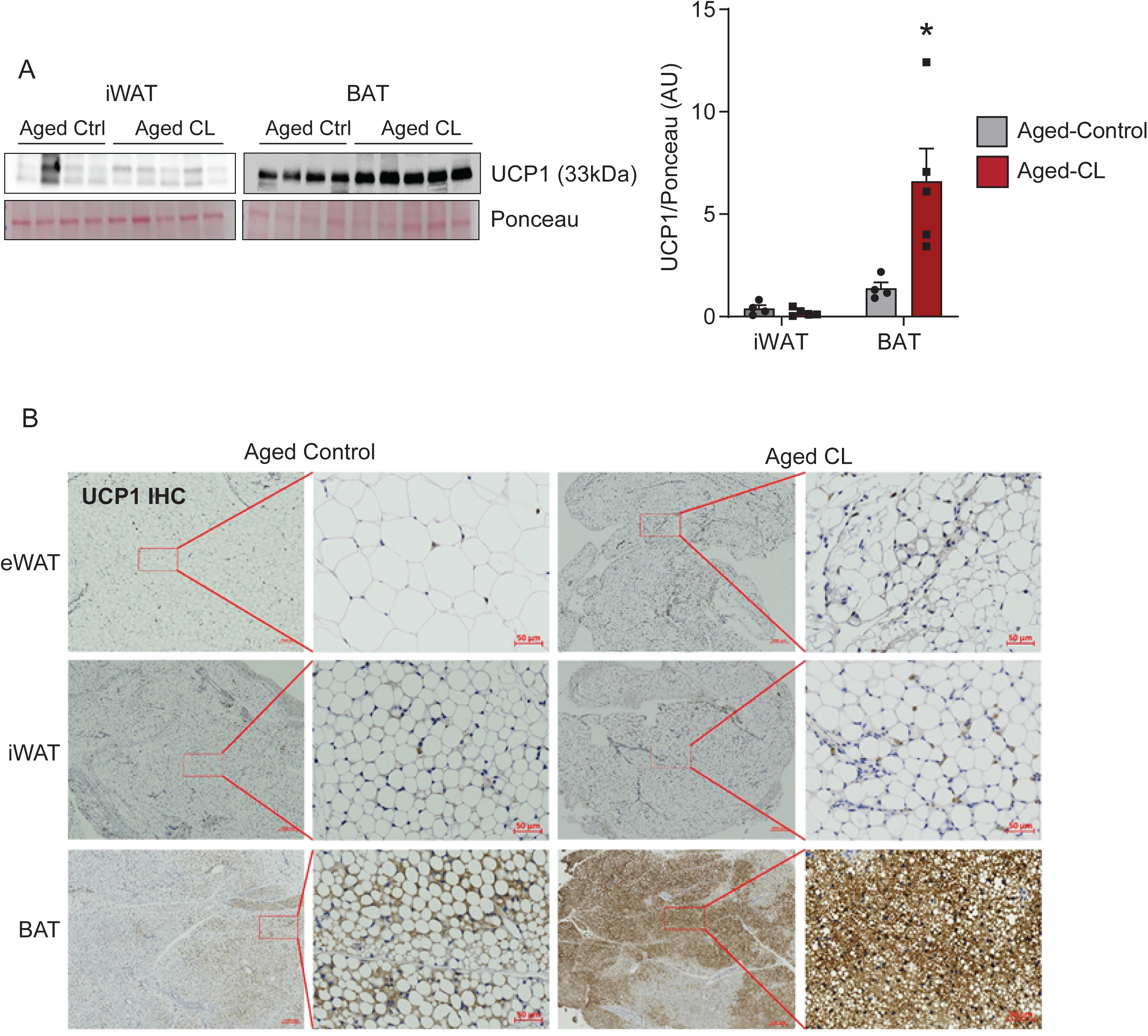
Effects of chronic β3-AR treatment on UCP1 expression in fat depots. (A) Western blot images and densitometry analysis for UCP1 in adipose depots (n = 4-5/group). (B) Representative images of UCP1 immunohistochemistry in adipose tissue sections of aged control and CL-treated mice. Please note that UCP1 expression (brown staining) was increased in the BAT but not in the eWAT and iWAT of CL-treated aged animals. Data are shown as average ± SEM; *p < 0.05 between the groups based on the Student’s-t-test analysis.

Increased metabolic activity in the absence of UCP1 in the WAT prompted us to investigate cellular mechanisms that could increase energy expenditure in a UCP1-independent manner. In particular, we focused on ATP-consuming TG-FA cycling, which has emerged as a major contributor to energy expenditure in UCP1 KO mice^19^. Futile lipid cycling could occur in two ways: lipolysis and subsequent re-esterification of released fatty acids, and lipolysis coupled with lipid oxidation and de novo lipogenesis (DNL)^19,36^. We assessed the protein expression of GYK, PEPCK, ACSL1, and DGAT1, the key enzymes involved in the aforementioned cycling mechanisms. GYK supports re-esterification by generating glycerol-3 phosphate (G-3-P) from glycerol, whereas PEPCK plays a key role in the de novo synthesis of G-3-P from sources other than glucose and glycerol. ACSL1 activates FAs for reesterification and DGAT1 catalyzes the final step of reesterification through the conversion of diacylglycerol and fatty acyl CoA to TG. In the iWAT, the expression of all these proteins was increased, however, in eWAT only GYK and PEPCK expression was upregulated with CL treatment in aged mice (Fig. 5A). Unlike WAT, the protein targets related to futile cycling were not impacted by CL treatment in the BAT of aged mice. To further validate these findings, we directly measured lipid turnover in the WAT depots using the ^2^H_2_O technique^19,20^. We measured ^2^H _2_O enrichment in the palmitate as a measure of DNL. On the other hand, ^2^H _2_O incorporation into glycerol indicates total TG turnover and glyceroneogenesis to support new TG synthesis in WAT. Both DNL and glycerol turnover expressed as fractional de novo synthesis (%) were significantly increased in both WAT depots with CL treatment in aged mice (Fig. 5B-C, top panels). Since CL treatment resulted in significant reductions in fat mass, we additionally calculated the de novo synthesis rates normalized to the total adipose mass per depot (Fig. 5B-C, bottom panels). DNL and glycerol turnover remained significantly increased only in iWAT after normalization to tissue weight. These results indicate that chronic β3-AR stimulation activates lipid turnover and DNL to increase energy expenditure in the iWAT of aged mice.

**Figure 5:**
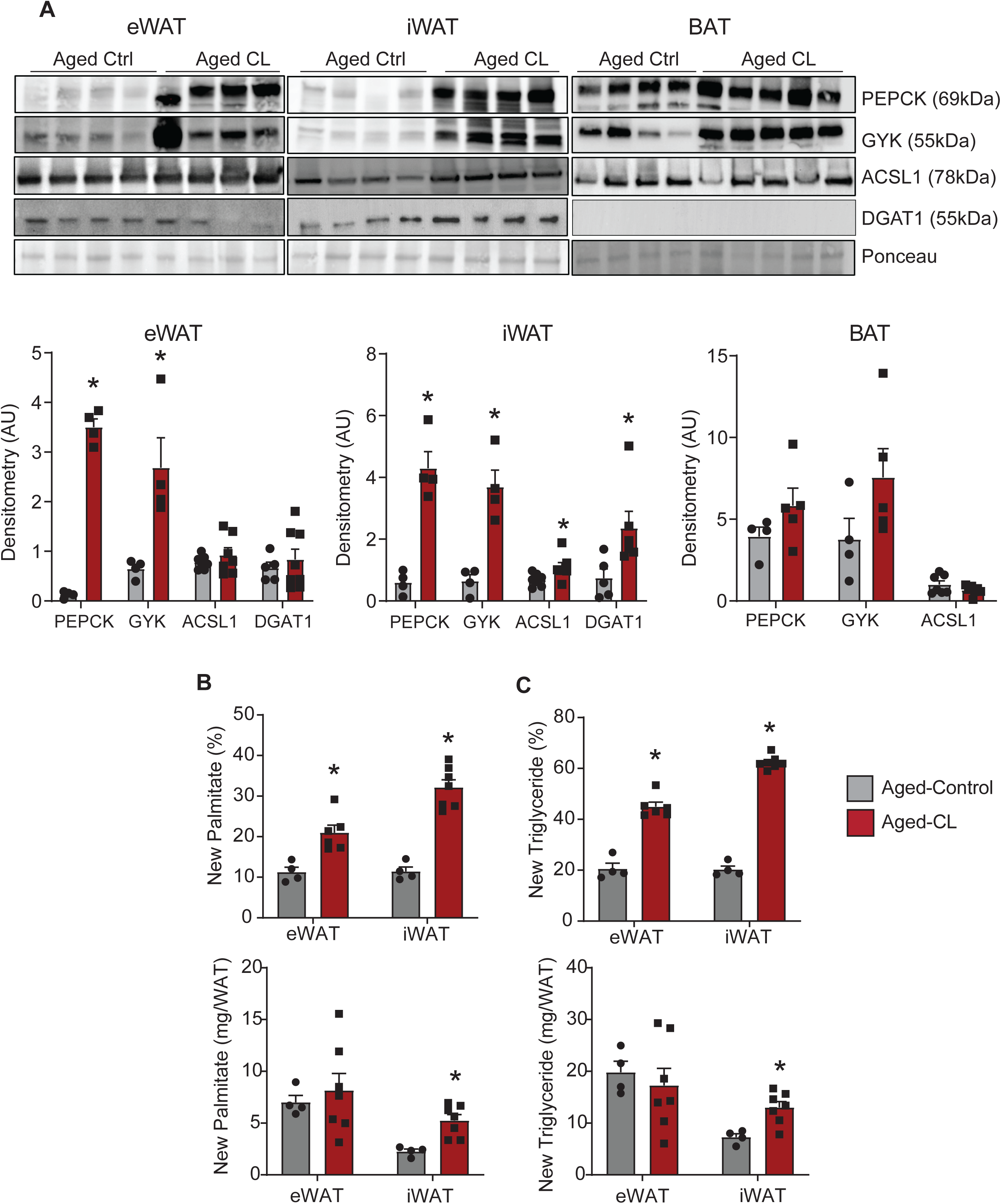
Effects of chronic β3-AR activation on lipid turnover and DNL in adipose tissues. A) Western blot images and densitometry analysis for quantitation of PEPCK, GYK, ACSL1, and DGAT1 protein levels in adipose depots (n = 4/group). (B) De novo synthesis of palmitate and C) glycerol in TG determined using the D_2_O technique (n=4-6/group). In both B and C, the top panels shows the data expressed as fractional de novo synthesis (%) and the bottom panel shows the enrichment de novo synthesis rates normalized to tissue weights. Data are shown as average ± SEM; *p < 0.05 between the groups based on the Student’s-t-test analysis.

### CL treatment elicits a dose-dependent inflammatory response in WAT

Aging increases inflammation both in the circulation and adipose tissue^37^. Hence, we investigated the impact of CL treatment on inflammatory markers, both in the serum and in different adipose depots. In eWAT, there was a trend for CL-induced decreases in IL6 and TNFα protein levels (Fig. 6A). However, we observed the opposite pattern in iWAT, where CL treatment increased the protein levels of IL6, IL10, and IL15, although MCP1 was significantly decreased (Fig. 6B). We did not observe any changes in cytokine or chemokine levels in the serum or BAT. The observed increase in inflammation in the iWAT (subcutaneous depot) could be due to insufficient expansion of the oxidative machinery, leading to a mismatch between FA supply and oxidation. We intended to test whether reducing the dose of CL (from 1 mg/kg to 0.5 mg/kg BW) would correct the imbalance in FA flux and elicit beneficial metabolic responses without inducing inflammation in the iWAT. Aged mice treated with a low dose of CL (0.5 mg/kg BW) displayed reduced fat mass, increased glucose tolerance, and increased mitochondrial activity in all three fat depots, as observed at the higher dose (Fig. 7A-D). Interestingly, unlike the higher dose, CL treatment @0.5 mg/kg reduced only the eWAT mass while preserving iWAT mass (Suppl. Fig. 2). More importantly, these metabolic improvements occurred without iWAT inflammation (Fig. 7E). Furthermore, low-dose CL treatment reduced circulating levels of IL6, an effect that was not observed at high doses (Fig. 7F).

**Figure 6:**
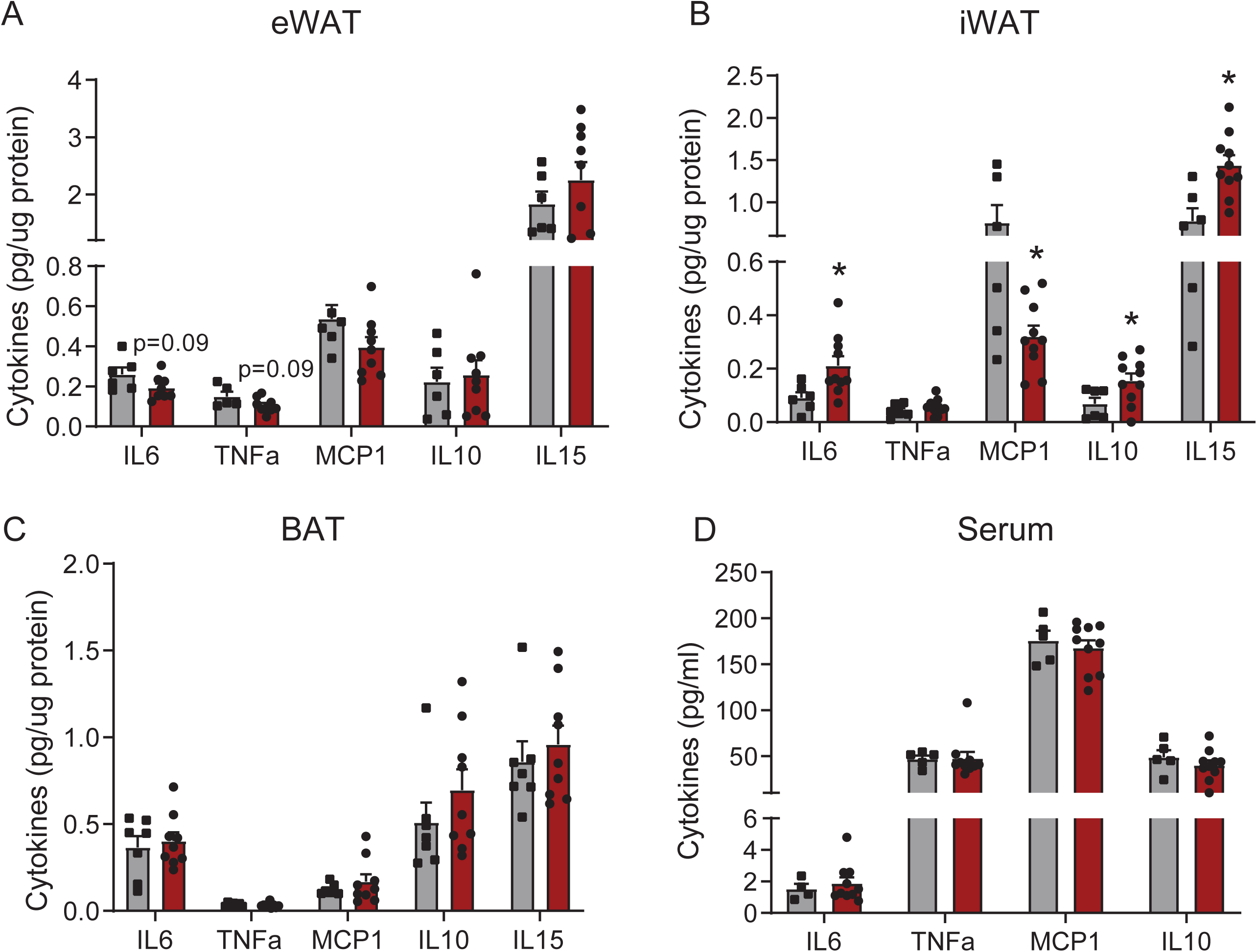
Changes in inflammatory mediators in serum and adipose tissue following CL treatment in aged mice. (A-D) Protein levels of inflammatory mediators were assessed in tissue lysates of eWAT, iWAT, and BAT, and serum samples of aged controls and CL-treated mice (n=6–8/group). Data are shown as average ± SEM; *p < 0.05 between the groups based on the Student’s t-test analysis.

**Figure 7:**
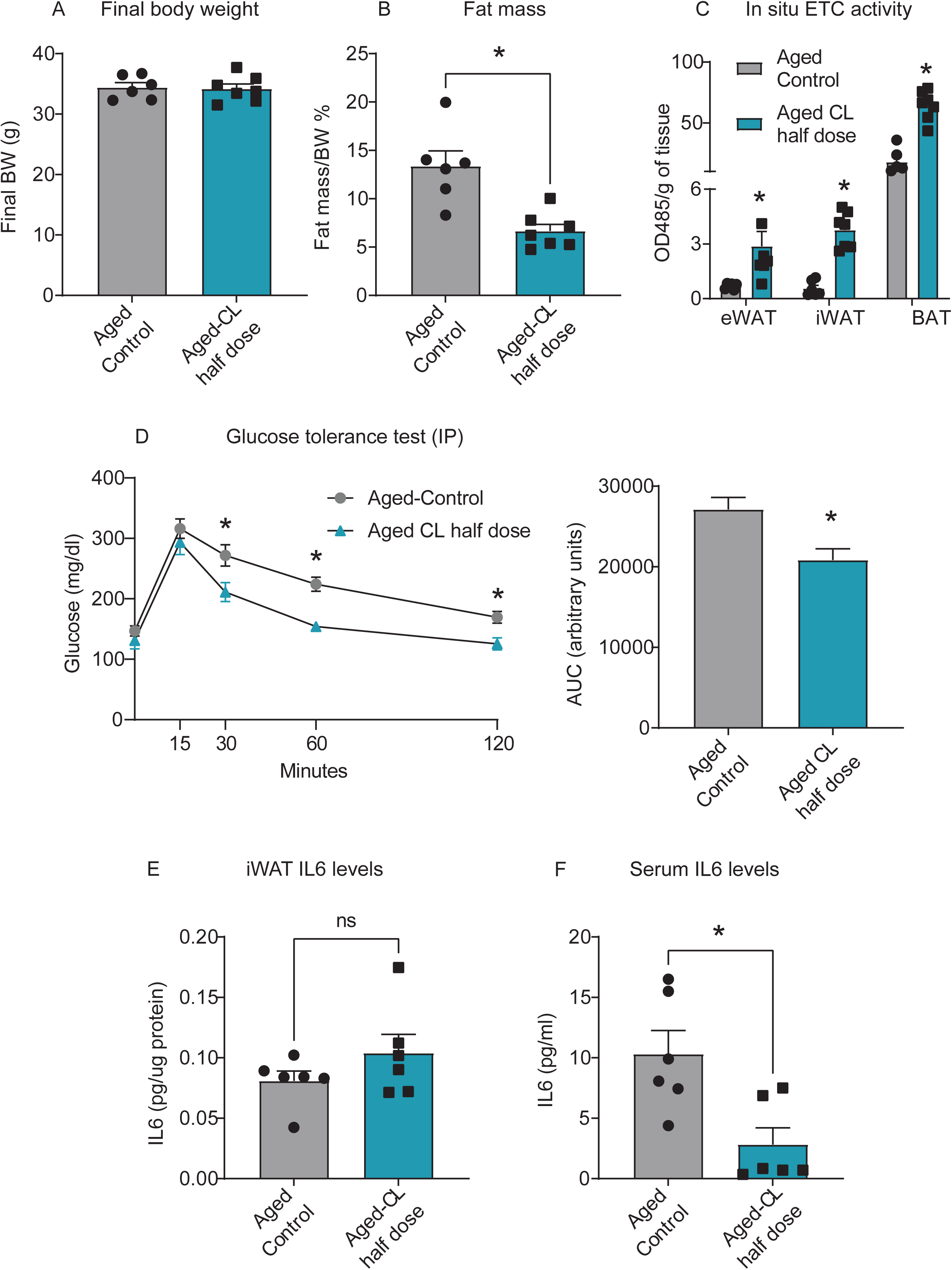
Effects of a reduced dose of CL on metabolic parameters and IL-6 levels in aged mice. Aged mice (18 months old; males) were treated with a reduced dose of CL 316,243 (0.5 mg/kg BW via osmotic pump infusion) for 4 weeks. (A) Final BW; (B) fat mass; (C) *In situ* ETC activity in adipose depots, and (D) Glucose values during intraperitoneal GTT and its AUC analysis. IL6 protein levels in (E) iWAT and (F) serum (n = 6–7/group). Data are shown as average ± SEM; *p < 0.05 between the groups based on the Student’s t-test analysis.

## Discussion

In the present study, we observed significant improvements in various metabolic parameters following chronic β3-AR treatment, including reduced fat mass, improved systemic glucose metabolism, increased energy expenditure, augmented oxidative capacity of adipose tissue, and elevated circulating adiponectin levels. At the cellular level, these favorable changes were likely mediated by UCP1-dependent thermogenesis in BAT and futile lipid cycling mediated increased energy expenditure in WAT. Moreover, a dose-dependent effect of CL treatment on inflammatory responses was evident, particularly in the iWAT depot.

UCP1-dependent thermogenesis predominantly occurs in classical brown and beige adipocytes, where heat is generated through the UCP1-mediated uncoupling of mitochondrial respiration from ATP synthesis. There are conflicting reports on UCP1 induction during aging, with findings varying depending on the method of thermogenic stimulation, age groups investigated, and duration of exposure^25,26,38,39^. We found that chronic β3-AR treatment significantly increased UCP1 expression and, in turn, the metabolic activity of BAT. This is consistent with a recent study by Tournissac et al.,^39^ who reported that CL treatment for a month increased BAT thermogenesis and glucose metabolism in 15-month-old mice. However, in line with previous studies^25,26^, beiging in response to β3-AR treatment was impaired with aging and we did not observe UCP1 expression in both WAT depots. This could be due to age-associated induction of senescence and the functional decline of adipose progenitor cells^40^, increased fibrosis^25^, and increased inflammation^41^. The increase in mitochondrial activity and fatty acid oxidation in the WAT depots without an increase in UCP1 expression suggested the possibility that alternate mechanisms of energy expenditure exist in WAT.

Activation of ATP-consuming futile cycling of metabolic substrates in beige adipocytes is a likely mechanism contributing to increased energy expenditure in the absence of UCP1. Specifically, we focused on TG-FA futile cycling which has recently emerged as a major contributor to cellular expenditure in UCP1-deficient mice^19^. Chronic β3-AR treatment increased the TG reesterification and glycerol and fatty acid turnover in the subcutaneous adipose depot (iWAT). This was accompanied by increased protein expression of GYK, PEPCK, ACSL1, and DGAT1 in the iWAT. It is estimated that 4 mol of ATP are expended for cycling 1 mol of TG (GYK and ACSL1 consume 1 and 3 ATP mol, respectively)^36,42^. During CL treatment, the increased demand for ATP synthesis required to support the futile lipid cycling drives glucose and FA mobilization and utilization leading to higher metabolic activity and energy expenditure in the iWAT. In addition to the direct impact on systemic energy homeostasis, recent evidence points to other putative functions of lipid cycling in regulating metabolic health. For example, Wunderling et al. reported that repeated lipolysis and reesterification will facilitate the regulation of the composition of stored FA pools by enabling FA modification and diversification^43^. In addition, lipokines derived from adipose DNL such as palmitoleate and fatty acid–hydroxy–fatty acids (FAHFAs) have been posited to positively impact systemic insulin sensitivity through endocrine actions^44,45^. However, it is yet to be determined whether activation of lipid cycling by itself is sufficient to improve systemic metabolism in aging.

With the high dose of CL (1mg/kg), the fatty acid influx from lipolysis likely overwhelmed the oxidative capacity, leading to inflammation. Correction of this mismatch through reducing the dose of CL (0.5 mg/kg BW) resulted in attenuation of inflammation in the iWAT. It is unclear why this effect was observed only in iWAT and not in eWAT. In addition, the fact that serum FFA and TG levels were unaltered, together with improved glucose tolerance, implies that these lipotoxic changes in the iWAT are local and contained. Hence, careful consideration should be given to the dose of lipolytic stimulation to maintain the metabolic benefits without inducing inflammation in the WAT.

Accumulating evidence supports the critical role of adipose tissue thermogenesis in longevity and delayed aging. Several genetic models and anti-aging interventions that delay aging and extend lifespan are associated with BAT activation and increased UCP1 expression in adipose depots^46–54^. These studies suggest that stimulation of adipose thermogenesis could have a beneficial impact on systemic aging and potentially delay the onset of age-related diseases. Supporting this notion, CL treatment has been shown to improve cognitive outcomes in 3xTg AD mice^39^, and mirabegron (an FDA approved β3-AR agonist) also improves pancreatic β**-**cell function, reduces lipotoxicity, and increases the number of type 1 fibers in the skeletal muscle of obese insulin-resistant middle-aged humans^16^. CL treatment-induced increase in adiponectin levels in our current study also aligns well with the idea of thermogenesis-mediated delayed aging, as higher circulating adiponectin levels are associated with longevity^55^.

In addition to thermogenesis, there is also an emerging role for lipolysis in health span regulation. Aging is associated with reduced lipolysis and reduced expression of proteins involved in lipolysis such as ATGL^56^. Further overexpression of ATGL-1 (homolog of ATGL) extends the lifespan in C. elegans while the reduction of ATGL-1 function suppresses the longevity of the long-lived mutants eat-2 and daf-2^57,58^. Similarly, overexpression of brummer (bmm, the major TG hydrolase in Drosophila) improved several healthspan outcomes like fertility, locomotion, mitochondrial function, and oxidative metabolism in Drosophila^59^. Our present studies support these previous findings in lower organisms and show that lipolytic stimulation using β3-AR agonists promotes metabolic health in aged mice.

Although these findings are promising, there are important caveats. First, our study design did not address the relative contribution of BAT thermogenesis (UCP1-dependent) and TG-FA cycling in WAT (UCP1-independent) to the observed beneficial systemic outcomes in response to adrenergic stimulation in aged mice. Future studies involving the removal of BAT before adrenergic stimulation will elucidate the sufficiency of WAT thermogenesis in correcting metabolic dysfunction during aging. Also, it is unclear why futile cycling is activated only in the subcutaneous and not in the visceral depot. Additionally, the cardiovascular side effects associated with chronic β3-AR agonist treatment must be addressed. Mirabegron induces a mild increase in systolic blood pressure at a dose that activates BAT in young participants^12^. Careful dose-response studies will identify the ideal dosage exerting beneficial metabolic effects in aged animals without inducing cardiovascular side effects through non-specific adrenergic receptor activation. It is also worth mentioning here that there have been recent developments in identifying novel agents that target non-β3-AR mechanisms to promote lipolysis (such as ABHD5 ligands^60^) which might offer safer clinical alternatives for the elderly population in the future. Finally, our current study design did not assess the site of palmitate synthesis (adipose tissue vs. other systemic tissues). However, the lack of TG accumulation in the liver and muscle with CL treatment (Suppl. Fig. 1G) suggests that fat tissue is likely a site of DNL in our study.

Overall, our findings underscore the promising potential of adipose tissue thermogenesis as a strategy to correct metabolic dysfunction and potentially delay systemic aging in older populations. More importantly, the preservation of futile TG-FA cycling mechanisms in aged WAT offers strategies that increase lipid turnover as a potential intervention for promoting healthy aging and combating age-related diseases.

## Supplementary

**Supplemental Fig. 1:**
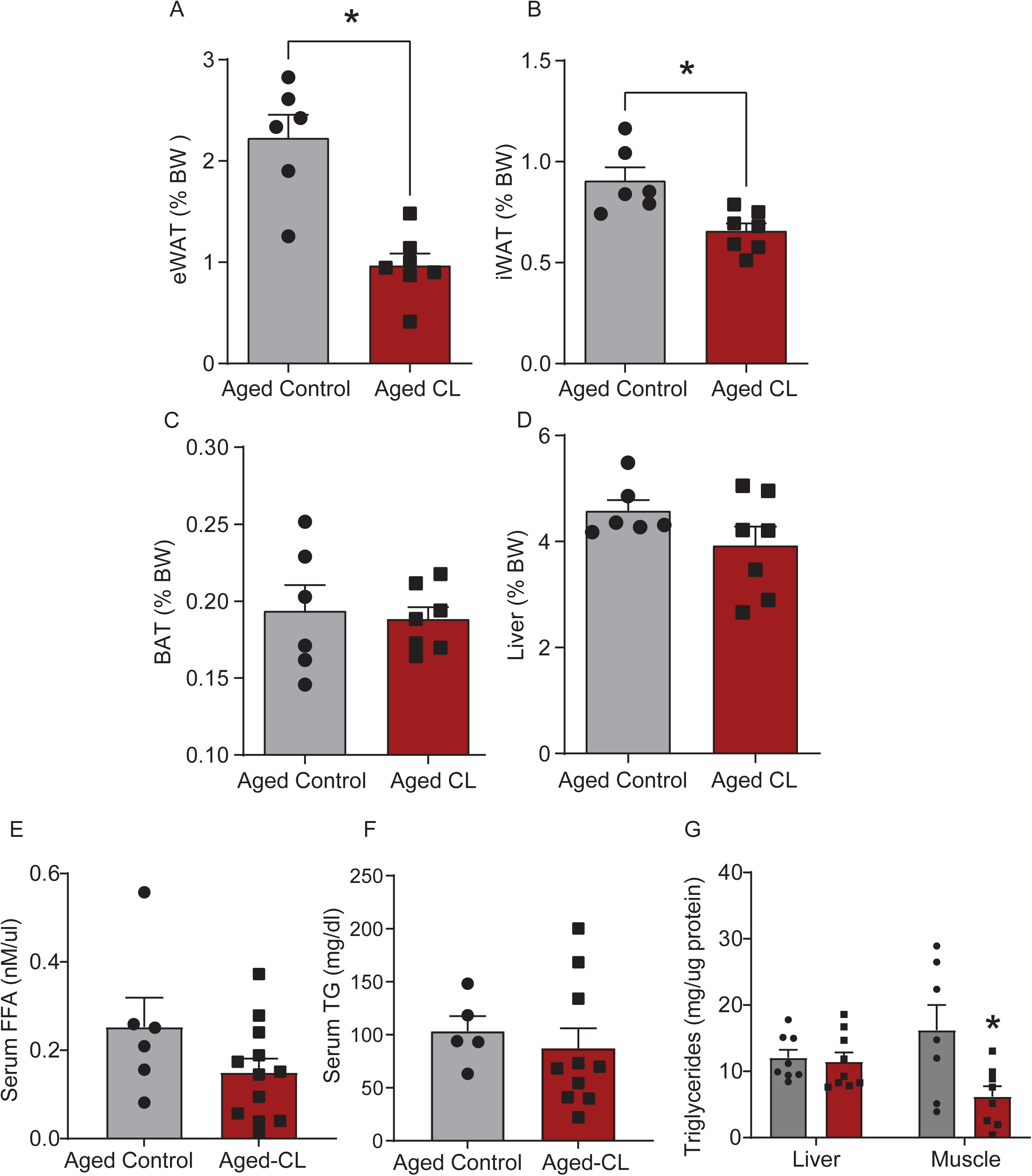
Effects of chronic β3-AR treatment on (A-C) individual adipose tissue and (D) liver weights, expressed as % BW, (E) serum FFA (F-G) serum and tissue TG levels (n = 6– 10/group). Data are shown as average ± SEM; *p < 0.05 between the groups based on the Student’s t-test analysis.

**Supplemental Fig. 2:**
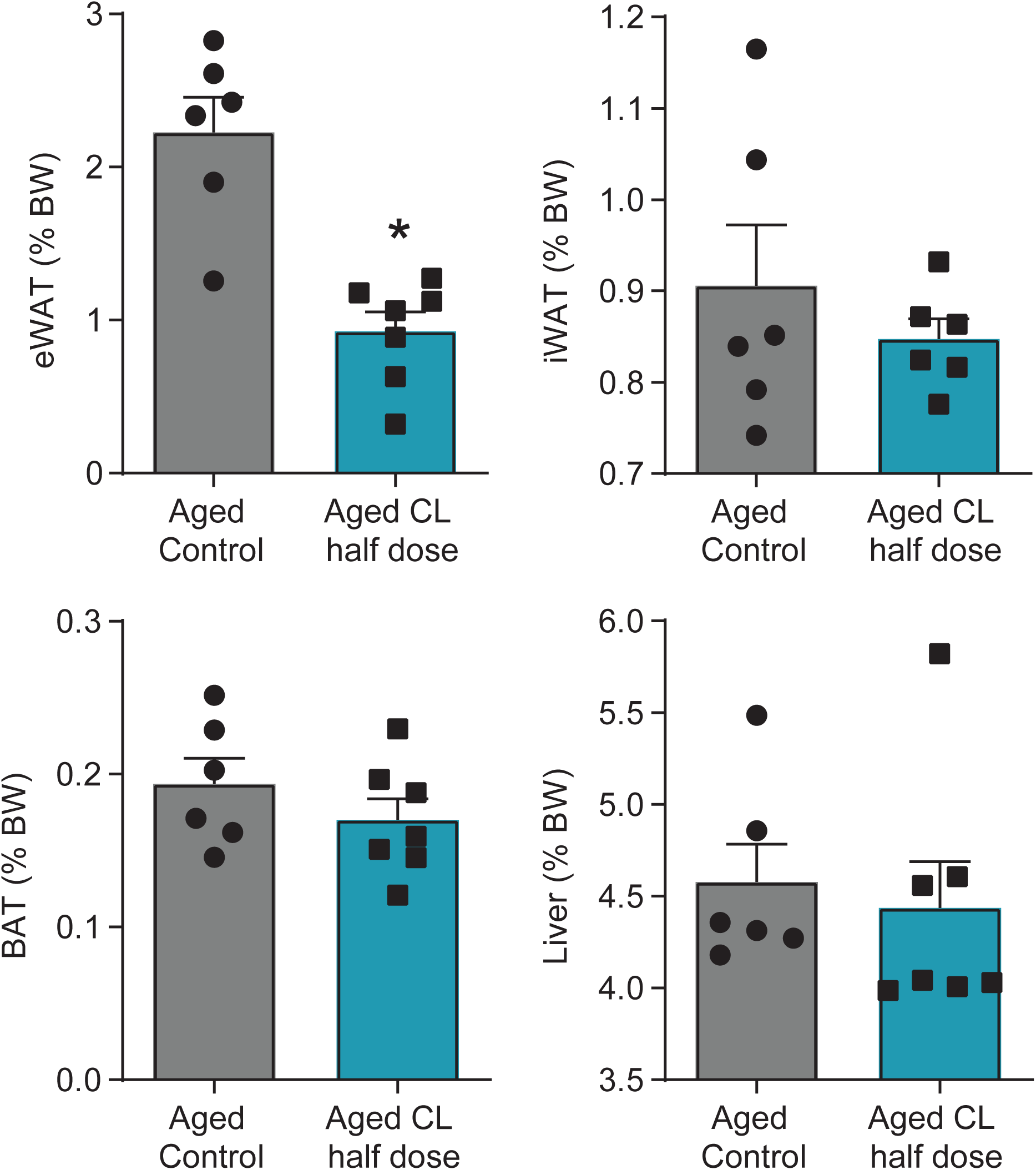
Individual tissue weights of aged control and CL-treated (0.5mg/kg) mice. (n = 6–7/group). Data are shown as average ± SEM; *p < 0.05 between the groups based on the Student’s t-test analysis.

## Acknowledgments

This work was supported by grants from the NIH (K01AG073613), American Heart Association (CDA1048544), Presbyterian Health Foundation, and OUHSC College of Medicine Alumni Association (COMAA) to PB, as well as from the National Institute on Aging (RF1AG072295, R01AG055395, R01AG068295, and R01AG070915) to AC and ZU. The histological services provided by the Stephenson Cancer Tissue Pathology Core were supported partly by National Institute of General Medical Sciences Grant P20GM103639 and National Cancer Institute Grant P30CA225520 from the National Institutes of Health.

## Author Contributions

Conceptualization, DN and PB; methodology and investigation, DN, BP, KT, SE, MR, WW, SC and MM; writing—original draft preparation, DN and PB; review and editing, DN, MS, ST, AY, MR, WW, MM, SC, ZU, AC and PB; funding acquisition, PB.

## Data Availability Statement

The data that support the findings of this study are available from the corresponding author upon reasonable request.

